# Therapeutic adenine base editing corrects nonsense mutation and improves visual function in a mouse model of Leber congenital amaurosis

**DOI:** 10.1101/2021.01.07.425822

**Authors:** Dong Hyun Jo, Hyeon-Ki Jang, Chang Sik Cho, Jun Hee Han, Gahee Ryu, Youngri Jung, Sangsu Bae, Jeong Hun Kim

## Abstract

Leber congenital amaurosis (LCA) is an inherited retinal degeneration that causes severe visual dysfunction in children and adolescents. In patients with LCA, pathogenic variants are evident in specific genes, such as *RPE65,* which are related to the functions of retinal pigment epithelium and photoreceptors. Base editing confers a way to correct pathogenic substitutions without double-stranded breaks in contrast to the original Cas9. In this study, we prepared dual adeno-associated virus vectors containing the split adenine base editors with *trans*-splicing intein (AAV-ABE) for *in vivo* adenine base editing in retinal degeneration 12 *(rd12)* mice, an animal model of LCA, which possess a nonsense mutation of C to T transition in the *Rpe65* gene (p.R44X). AAV-ABE induced an A to G transition in retinal pigment epithelial cells of *rd12* mice when injected into the subretinal space. The on-target editing was sufficient to recover wild-type mRNA, RPE65 protein, and light-induced electrical responses of retinal tissues. We suggest adenine base editing to correct pathogenic variants in the treatment of LCA.

## Introduction

Biallelic mutations in the *RPE65* gene as well as other genes related to the functions of the retinal pigment epithelium (RPE) and photoreceptors cause Leber congenital amaurosis (LCA), an inherited retinal degeneration (1, 2). Patients with LCA typically show severe early-onset visual dysfunction (3). Until the approval of the first-in-class adeno-associated virus (AAV)-based gene therapy (voretigene neparvovec; Luxturna, Spark Therapeutics), there were no cures for LCA. In addition to AAV delivering the *RPE65* gene via subretinal injection (4), genome editing by clustered regularly interspaced short palindromic repeats (CRISPR) can provide alternative modalities to treat patients with LCA by direct modification of pathogenic variants in the genome (5). It is noteworthy that genome editing tools permanently corrects the pathogenic mutations at the endogenous locus. Small insertions and deletions (indels) after DNA double-stranded breaks (DSBs) can correct a frame-shift intron variant of the *CEP290* gene (IVS26; c.2991+1655 A>G), the most frequent one in patients with LCA type 10 (6). On the other hand, we have shown the therapeutic potential of homology-directed repair (HDR) in the previous study (7). Dual AAVs carrying CRISPR-Cas9 and donor DNA in each vector resulted in ~1.2% of HDR and ~1.6% of in-frame 1-codon deletion when they were administered through subretinal injection in retinal degeneration 12 *(rd12)* mice (7).

Although HDR is a method of gene recovery by CRISPR (8), the correction efficiencies seemed insufficient to guarantee clinical efficacy (7). Besides, there are concerns that CRISPR-mediated DSBs frequently induce unexpected large deletion and complex rearrangement (9, 10). In contrast, base editing is a more direct tool to correct and recover disease phenotypes (11). In particular, adenine base editors (ABEs) that consist of the evolved version of the tRNA-specific adenosine deaminase (TadA) fused to nuclease-deficient Cas9 convert A to G without DSBs (12).

About 75% of likely pathogenic/pathogenic variants of the *RPE65* gene, one of the most frequently mutated gene in patients with LCA (13, 14), are substitutions in the ClinVar database. Similar to patients with LCA type 2, *rd12* mice possess a nonsense mutation of C to T transition at position 130 in the exon 3 of the *Rpe65* gene (p.R44X) (15). It is noteworthy that five independent groups have reported that this R44X mutation is evident in humans (http://www.ncbi.nlm.nih.gov/clinvar/variation/374497) (14, 16, 17). Adenine base editing provides a practical approach to induce the intended transition to recover wild-type sequence with higher on-target activities and negligible small indels that are derived after DNA DSBs by CRISPR effectors (18).

Here, we demonstrated the therapeutic potential of adenine base conversion of a nonsense mutation in the *Rpe65* gene of *rd12* mice. Adenine base editors (ABEs) subretinally delivered via dual AAV serotype 9 vectors effectively induced an A to G transition in the RPE *in vivo.* This therapeutic base editing of the *Rpe65* gene increased light-induced electrical responses of the retinal tissues after dark adaptation in *rd12* mice which exhibit no responses without treatment, suggesting a therapeutic opportunity of base editing in the treatment of patients with LCA.

## Results

### Adenine base editing to correct a nonsense mutation in rd12 mice

To correct a nonsense mutation of the *Rpe65* gene in *rd12* mice, we utilized an optimized version of ABE, ABEmax (19), employing the protospacer adjacent motif (PAM) of NG sequences instead of original NGG (20), named NG-ABEmax (21). In this approach, the disease-associated point mutation (c.130C>T) was at the sixth position from the 5’ end of the protospacer (A6) (**Figure 1A**). As of now, AAV is the most reliable and effective way to deliver ABEs *in vivo* (22). It is also noteworthy that AAV-based gene therapy is a clinical approach approved by the Food and Drug Administration of the United States to treat patients with LCA. To bypass the size limit of the cargo of AAVs (~4.8 kb), we divided the NG-ABEmax into halves using a *trans*-splicing intein (23, 24) (**Figure 1B**). We then tested whether each half of NG-ABEmax encoded by plasmids would exhibit an editing activity via intein-mediated reconstitution. When we treated embryonic fibroblasts from *rd12* mice with dual plasmids encoding each half of NG-ABEmax, we observed 5.4±0.7% of editing efficiency, showing the validity of our split NG-ABEmax system (**Supplemental Figure 1**). Based on our previous experiments of AAV-mediated *in vivo* genome editing (7), we decided to test the therapeutic efficacy of ABEs at 6 weeks after the subretinal injection (**Figure 1C**).

**Figure 1.**
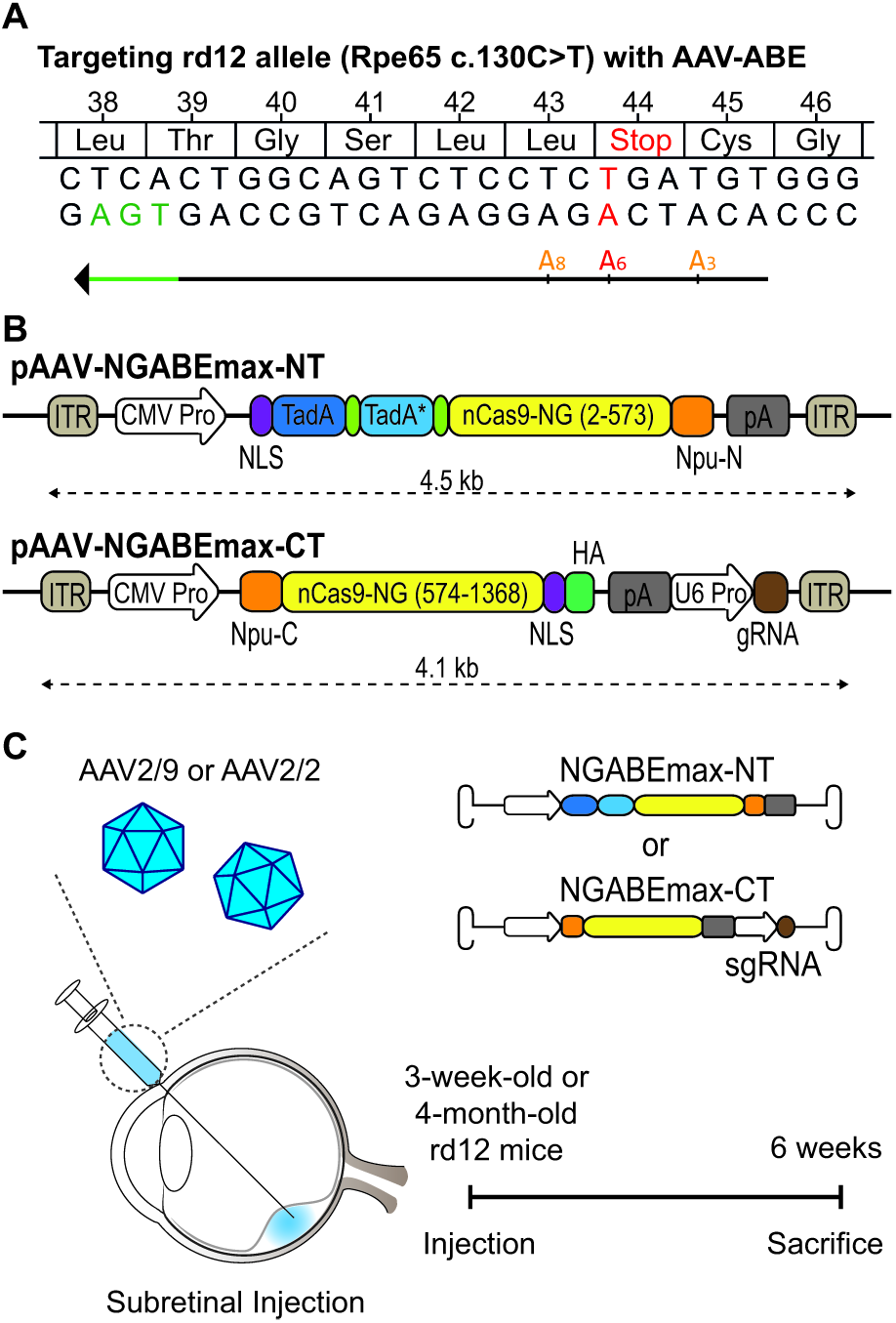
Adenine base editing to correct a nonsense mutation in *rd12* mice. **(A)** The C>T nonsense mutation (red) at position 130 (c.130C>T; p.R44X). Using NG PAM (TGA, green) ABEmax, the targeted adenosine is located at protospacer position 6, counting the PAM as positions 21-23. The other adenosine within the editing window of ABEmax (A3 and A8) were indicated in yellow-orange letters. **(B)** A schematic drawing of the dual AAV vectors for ABE delivery. CMV Pro, CMV promoter; gRNA, guide RNA; ITR, inverted terminal repeat; nCas9-NG, NG PAM ABEmax; NLS, nucleus localization signal; Npu-C, C-intein from *Nostoc punctiforme*; Npu-N, N-intein from *Nostoc punctiforme*; pA, polyA; TadA, tRNA-specific adenosine deaminase; TadA*, the evolved version of TadA; U6 Pro, U6 promoter. **(C)** A schematic diagram depicting the design of animal experiments. The dual AAV vectors were packaged into AAV serotype 2 or 9. 3-week-old or 4-month-old *rd12* mice received subretinal injection of AAV-ABE. Six weeks later, the eyes were enucleated for further analyses.

### In vivo adenine base editing based on AAV corrects a nonsense mutation in the RPE of rd12 mice

Although almost all the tested serotypes of AAVs transduced the RPE cells through the subretinal injection (25), there is no direct evidence on which serotypes are better in delivering dual AAV vectors for adenine base editing of RPE cells. A notable finding is that AAV serotype 9 effectively induced HDR in our previous study (~1.2%) (7), whereas AAV serotype 1 did not in a study by Suh et al. (0.03%) (26). To select the most reliable AAV serotype for the ABE delivery in RPE cells, we tested AAV serotype 9 and serotype 2 which is utilized in voretigene neparvovec, a gene therapy drug for LCA in 4-month-old *rd12* mice. Strikingly, we observed a higher on-target editing (A6 to G) efficiency with AAV serotype 9 (13.5±0.3%) than serotype 2 (0.3±0.2%) (**Figure 2, A and B**). When the target adenosine is at A6, the other adenosines (A3 and A8) lay within the activity window of ABEmax, about 3 to 8 positions from the 5’ end of the protospacer, counting the PAM (TGA) as positions 21-23 (19). In total, the anticipated products of bystander editing at A1, 3, 6, or 11 are C45C (synonymous), C45R, L43P, or L42P (non-synonymous), respectively. Among total base-edited reads at any sites, the top 5 products were X44R (the desired edit, 67.5%), X44R and L43P (17.8%), L43P (4.2%), X44R and C45R (4.1%), and C45R (2.2%) (**Figure 2C**). It is noteworthy that L43P and C45R variants have not been reported in patients with inherited retinal degeneration.

**Figure 2.**
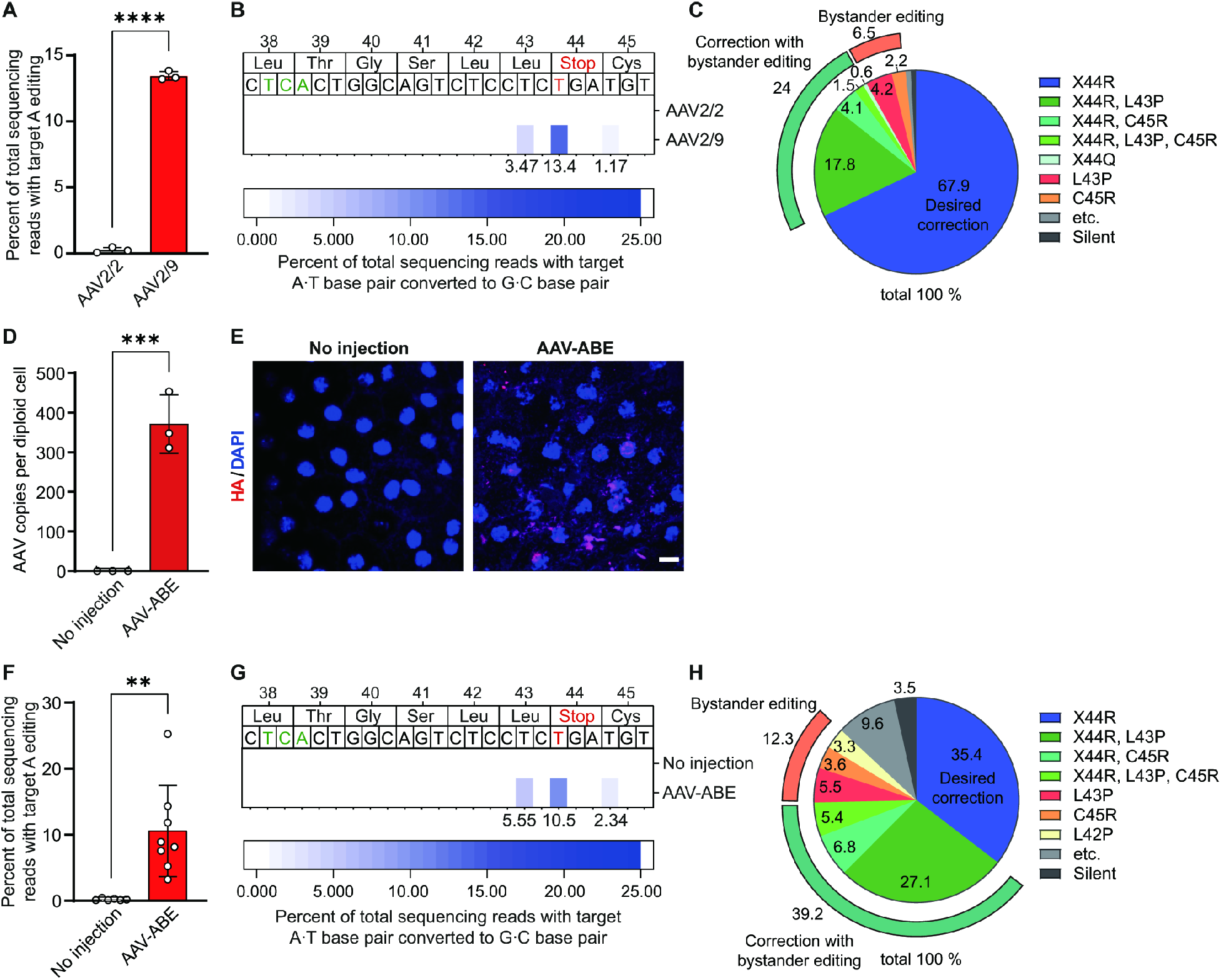
*In vivo* adenine base editing based on AAV corrects a nonsense mutation in the RPE of *rd12* mice. **(A)** Quantitation of the intended A to G correction at position 6 (A6) in the RPE cells at 6 weeks after the subretinal injection of AAV serotype 2 (AAV2/2) or serotype 9 (AAV2/9) containing NG-ABEmax on 4-month-old *rd12* mice *(n* = 3). \*\*\*\**P* < 0.0001, two-tailed Student’s t-test. **(B)** The heatmap of DNA reads according to the edited nucleotide positions within the *Rpe65*. **(C)** The editing outcomes including the intended (X44R) and bystander edits in the RPE cells at 6 weeks after the subretinal injection of AAV serotype 9 containing NG-ABEmax on 4-month-old *rd12* mice *(n* = 3). **(D)** Quantitation of AAV copies per diploid cell of the RPE *(n* = 3). \*\*\**P* < 0.001, two-tailed Student’s t-test. **(E)** Representative photographs of confocal microscopy of the RPE of *rd12* mice at 6 weeks after the subretinal injection of AAV-ABE. Scale bar, 10 μm. **(F)** Quantitation of the intended A to G correction at position 6 (A6) in the RPE cells at 6 weeks after the subretinal injection of AAV serotype 9 containing NG-ABEmax on 3-week-old *rd12* mice *(n* = 6 for ‘No injection’ group and *n* = 8 for ‘AAV-ABE’ group). \*\**P* < 0.01, two-tailed Student’s t-test. **(G)** The heatmap of DNA reads according to the edited nucleotide positions within the *Rpe65*. **(H)** The editing outcomes including the intended (X44R) and bystander edits in the RPE cells at 6 weeks after the subretinal injection of AAV serotype 9 containing NG-ABEmax on 3-week-old *rd12* mice *(n* = 8).

In lines with *in vivo* adenine base editing, we observed multiple AAV copies in the RPE cells of *rd12* mice at 6 weeks after the subretinal injection of dual AAV serotype 9 vectors containing NG-ABEmax (AAV-ABE) (**Figure 2D**). Furthermore, AAV-mediated ABE delivery was supported by the prominent nuclear and cytoplasmic expression of hemagglutinin (HA) tag (**Figure 2E**) which is encoded by the sequence in the AAV vectors (**Figure 1B**). Promoted by the effective delivery of ABEs and resultant adenine base editing, we then injected AAV-ABE into the subretinal space of 3-week-old *rd12* mice. As in 4-month-old mice, AAV-ABE effectively induced on-target editing of 10.6±6.9% (range, 3.2 to 25.3%) (**Figure 2, F and G**). The top 5 products among base-edited reads in 3-week-old mice were X44R (35.2%), X44R and L43P (27.1%), X44R and C45R (6.8%), L43P (5.5%), followed by X44R, C45R, and L43P (5.4%) (**Figure 2H**).

### In vivo adenine base editing recovers RPE65 protein in the RPE and improves the visual function of rd12 mice

In treating patients with LCA possessing the pathogenic variants in the *RPE65* gene, the purpose of both gene therapy and genome editing is restoring the functional RPE65 protein, which is essential in the visual cycle (1, 27). mRNA sequencing analyses showed that most (89.3±2.3%) of the reads of the *Rpe65* mRNA from the RPE tissues of AAV-ABE-treated mice were the products of A6 correction (**Figure 3A**), which demonstrated a much higher level of correction than expected from the analysis of DNA correction efficiency (**Figure 2F**). As expected from the targeted deep sequencing, the top 2 reads of the *Rpe65* mRNA corresponded to X44R (24.9%) and X44R and L43P (23.9%), whereas non-corrected mRNA reads were only 2.0 % of total reads. (**Figure 3, A and B and Supplemental Figure 2**). Due to a nonsense mutation of R44X in the *Rpe65* gene, *rd12* mice lack the RPE65 protein (**Figure 3C**) (15, 26). The RPE cells in the AAV-ABE-treated *rd12* mice expressed the RPE65 protein with the prominent expression of HA tag in the nucleus (**Figure 3C**). To investigate whether the restoration of the RPE65 protein by AAV-ABE led to functional recovery, we performed electroretinography (ERG) with wild-type C57BL/6 and *rd12* mice after dark adaptation. As in the previous study employing lentiviral vectors for ABE delivery (26), AAV-ABE increased light-induced electrical responses of retinal tissues in *rd12* mice (**Figure 3D**). The amplitudes of a- and b-waves of dark-adapted ERG responses of AAV-ABE-treated *rd12* mice at 0 dB light stimuli were 30.4±3.4% and 29.2±2.5% of those of wild-type mice, respectively (**Figure 3, E and F**). We further measured the optokinetic responses to respond to the rotating stimuli in a virtual cylinder. In this experiment estimating the visual function, AAV-ABE-treated *rd12* mice significantly recovered the visual thresholds compared to untreated ones (**Figure 3G**).

**Figure 3.**
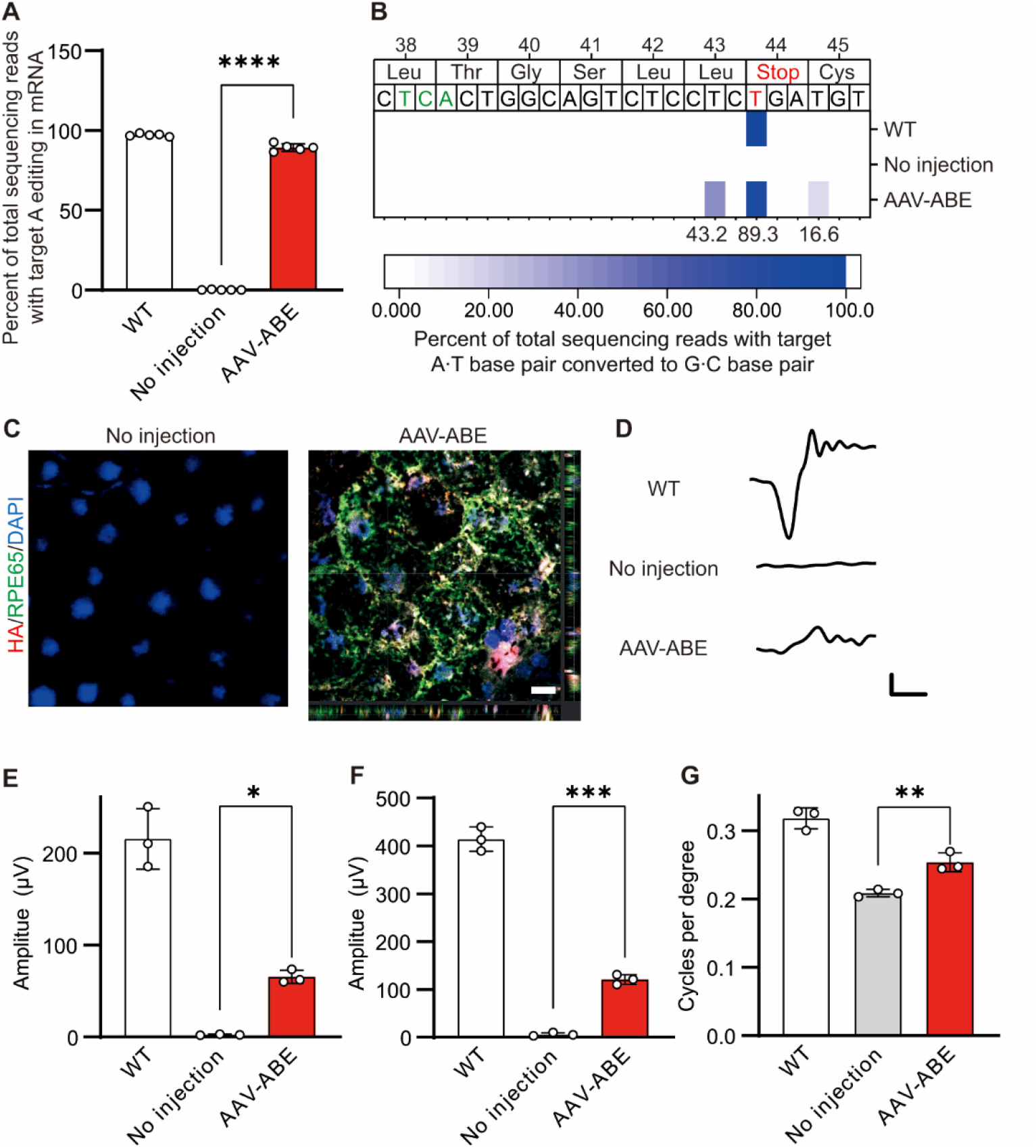
*In vivo* adenine base editing recovers RPE65 protein in the RPE and improves visual function of *rd12* mice. **(A)** Quantitation of the A6-corrected mRNA in sequencing of the *Rpe65* mRNA of the RPE cells from C57BL/6 and 3-week-old *rd12* mice according to the treatment of AAV-ABE (*n* = 5). \*\*\*\**P* < 0.0001, one-way ANOVA with post-hoc Tukey’s multiple comparison tests. WT, wild-type C57BL/6. **(B)** The heatmap of mRNA reads according to the edited nucleotide positions within the *Rpe65* mRNA. WT, wild-type C57BL/6. **(C)** Representative photographs of confocal microscopy of the RPE at 6 weeks after the subretinal injection of AAV-ABE on 3-week-old *rd12* mice. Scale bar, 10 μm. **(D)** Representative waveforms of ERG of wild-type C57BL/6 and *rd12* mice according to the treatment of AAV-ABE. WT, wild-type C57BL/6. **(E)** Quantitative analyses of amplitudes of a-waves of ERG of C57BL/6 and *rd12* mice according to the treatment of AAV-ABE (*n* = 3). \**P* < 0.05, one-way ANOVA with post-hoc Tukey’s multiple comparison tests. WT, wild-type C57BL/6. **(F)** Quantitative analyses of amplitudes of b-waves of ERG of wild-type C57BL/6 and *rd12* mice according to the treatment of AAV-ABE (*n* = 3). \*\*\**P* < 0.001, one-way ANOVA with post-hoc Tukey’s multiple comparison tests. WT, wild-type C57BL/6. **(G)** Quantitative analyses of the visual acuity of wild-type C57BL/6 and *rd12* mice according to the treatment of AAV-ABE (*n* = 3). \*\**P* < 0.01, one-way ANOVA with post-hoc Tukey’s multiple comparison tests. WT, wild-type C57BL/6.

### Analysis of potential off-target effects of in vivo adenine base editing

Although ABEs do not induce DSBs, there are still concerns of single-guide RNA (sgRNA)-dependent off-targeting of DNA (A to G conversion at other sites in the genome) (28) and sgRNA-independent RNA deamination (29–31) by an excess wild-type TadA protein included in ABEmax. Despite effective on-target editing (13.3±8.9%), we only observed negligible off-target editing of DNA at the 2 predicted sites from the Cas-OFFinder (32) and the 10 predicted sites from the Circle-seq (26, 33) in the RPE samples of AAV-ABE-treated *rd12* mice (**Figure 4A and Supplemental Table 1**). However, we observed one meaningful RNA deamination patterns at MCM3AP transcript among the 5 tested sites (29–31), which were predicted on the sequence similarity to the TadA substrate (GCUCGGCUACGAACCGAG) (34) (**Figure 4B and Supplemental Table 2**). Although the mRNA mutations are transient in cells, it is necessary to use an advanced version of ABEs, such as ABE8eW (35), ABE8eWQ (36), and ABE8s with V106W (37) that show higher base editing activity and negligible RNA deamination effects in the future. For the clinical application, it is essential to minimize the toxicity to the adjacent tissues. Remarkably, AAV-ABE did not affect histologic integrity (**Figure 4C and Supplemental Table 3**) and ERG responses in wild-type C57BL/6 (**Figure 4, D and E**). These data implied that AAV-ABE did not induce significant nonspecific tissue damage upon local administration.

**Fig. 4.**
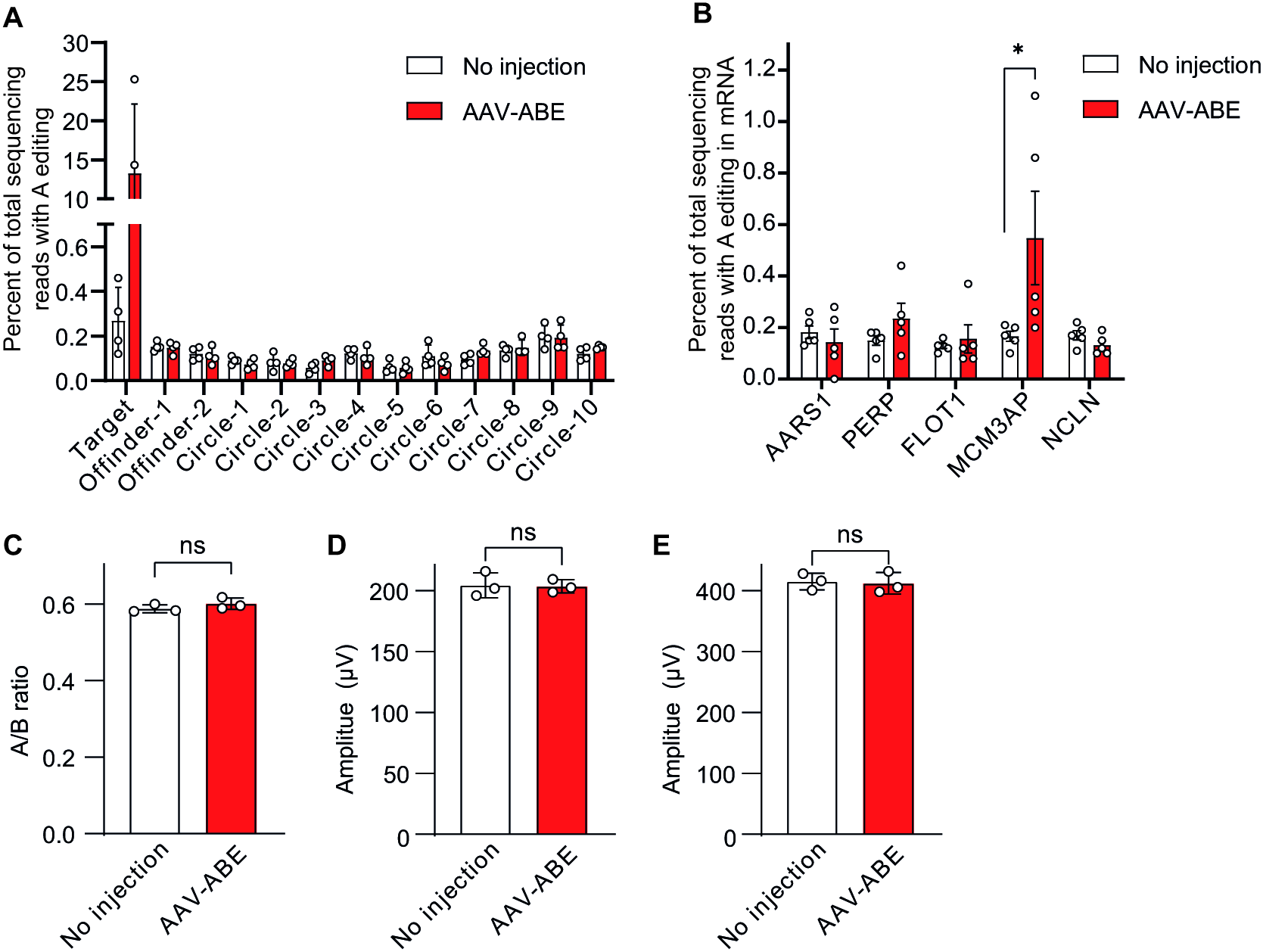
Analysis of potential off-target effects of *in vivo* adenine base editing. **(A)** On- and off-target editing rates of specific sites of the genomic DNA of the RPE tissues at 6 weeks after the after the subretinal injection of AAV-ABE on 3-week-old *rd12* mice (*n* = 4). Off-target sites were predicted from Cas-OFFinder (2 sites) and CIRCLE-seq (6 sites). **(B)** Off-target editing rates of RNA of the RPE tissues at 6 weeks after the subretinal injection of AAV-ABE on 3-week-old *rd12* mice (*n* = 5). *, *P* < 0.05; Mann-Whitney U test. Three off-target sites were selected based on the sequence similarity to the TadA substrate **(C)** Quantitation of the histologic integrity at 6 weeks after the after the subretinal injection of AAV-ABE on C57BL/6 mice (*n* = 3). *ns, P* > 0.05; two-tailed Student’s t-test. **(D)** Quantitative analyses of amplitudes of b-waves of ERG at 6 weeks after the after the subretinal injection of AAV-ABE on C57BL/6 mice (*n* = 3). A/B ratio indicates the ratio of the thickness from the internal limiting membrane to the inner nuclear layer (‘A’) to that from the internal limiting membrane to the outer nuclear layer (‘B’). *ns*, *P* > 0.05; two-tailed Student’s t-test. **(E)** Quantitative analyses of amplitudes of b-waves of ERG at 6 weeks after the after the subretinal injection of AAV-ABE on C57BL/6 mice (*n* = 3). *ns*, *P* > 0.05; two-tailed Student’s t-test.

## Discussion

In this study, we demonstrated that adenine base editing could correct a nonsense mutation and restore visual function in *rd12* mice, an animal model of LCA. Notably, we achieved ~10% of editing efficiencies in the intended correction with clinically applicable AAV vectors, leaving aside the additional ~3% of read-through edits (38). Considering we only injected AAV vectors into the subretinal space of less than half of the murine retina, we speculated that these results in mice could be improved in humans (7). In patients with LCA, retinal surgeons injected AAV vectors in the selected area near the macula to enhance the possibility of functional recovery of the central vision (4). On the other hand, in the experiments using mice, we measured editing rates in the whole RPE, although only less than half of RPE cells near the subretinal bleb were transduced by AAV When normalized to the transduction efficiency, we achieved enough editing efficiency to have clinical potential, as we demonstrated improved visual function in a mouse model.

It is also remarkable that we utilized clinically applicable AAV vectors for *in vivo* adenine base editing. Recently, Suh et al. reported that lentiviral vector-delivered ABE targeting NAG PAM enables effective restoration of visual function in adult *rd12* mice, similar to our results. As the authors mentioned in the paper, lentiviral vectors are only for proof-of-concept studies in the current form (26). In addition to the delivery system (clinically applicable AAV vs. lentivirus), we used the different ABE construct and sgRNA sequence, enabling comparable on-target editing to the lentivirus-based approach.

As in our previous study employing HDR and in-frame deletion by CRISPR (7), we observed higher mRNA (**Figure 3, A and B**) and functional recovery (**Figure 3, D-F**) than expected from DNA editing (~10%). ~90% of the total reads of the *Rpe65* mRNA from the RPE tissues of AAV-ABE-treated *rd12* mice were the products of the corrected DNA sequence. Furthermore, the amplitudes of ERG waves in the treated *rd12* mice were ~30% to those in wild-type C57BL/6 mice. We speculated that these phenomena are related to nonsense-mediated decay of mutant mRNA from the DNA sequence with a nonsense mutation (39). Permanent correction in the DNA can result in augmented clinical responses at the levels of mRNA, protein, and functions.

Taken together, we suggest an AAV-based adenine base editing as a permanent and direct therapeutic approach to treat patients with LCA-associated substitutions. The impact of base editing might be higher with the improved versions of base editors in terms of specificity and efficacy (35–37). Nonetheless, AAV-ABE in the current form effectively corrected a pathogenic variant and improved visual function in *rd12* mice. We expect this AAV-based adenine base editing to be applied to patients with LCA who only have limited treatment options.

## Methods

### Molecular cloning and virus production

All plasmids were constructed using Gibson-cloning with the NEBuilder HiFi DNA Assembly master mix (NEB). For the construction of the N-terminal part of the split NG-ABEmax-encoding vector (AAV-NT-ABEmax), the N-terminal coding sequence of ABEmax along with the CMV promoter (pCMV-ABEmax; Plasmid #112095, addgene) and N-intein from *Nostoc punctiforme* was PCR-amplified with matching overlaps and cloned into an AAV2-ITR backbone (NotI and XbaI restriction digest of pAAV-nEF-Cas9 plasmid; Plasmid #87115, addgene). For the construction of the C-terminal part of the split NG-ABEmax-encoding vector (AAV-CT-ABEmax), C-Intein from *Nostoc punctiforme* along with the CMV promoter, the C-terminal coding sequence of ABEmax and gRNA along with U6 promoter were PCR-amplified with matching overlaps and cloned into an AAV2-ITR backbone. Recombinant AAV packaging (AAV2/2 and AAV2/9) was performed by Vigene Biosciences.

### Animals

C57BL/6 mice and mating pairs of *rd12* mice (stock no. 005379, The Jackson Laboratory) were purchased from Central Laboratory Animal and maintained under a 12-hour dark-light cycle. *rd12* mice were mated each other to produce offspring. Three-week-old and four-month-old *rd12* mice underwent subretinal injection.

### Subretinal injection of AAV

After deep anesthesia, AAV-NT-ABEmax and AAV-CT-ABEmax (5.4 × 10^10^ vg for AAV2/2 and 7.3 × 10^10^ vg for AAV2/9 each in 3 μl PBS) were injected into the subretinal space of mice using a customized Nanofil syringe with a 33 G blunt needle (World Precision Instrument) under an operating microscope (Leica), as previously described (40).

### Targeted deep sequencing

Genomic DNAs and total RNAs were extracted from retinal or RPE tissues at 6 weeks after subretinal injection for the sequencing of DNA or RNA at on-target or off-target sites. Each tissue sample was sonicated for a few seconds with the buffer LBP1 from NucleoSpin RNA Plus kits (MACHEREY-NAGEL) and divided into two tubes. The lysates were then purified to genomic DNA and RNA using NucleoSpin Tissue Kits and NucleoSpin RNA Plus kits (MACHEREY-NAGEL), respectively, according to the manufacturer’s instructions. The purified RNA (total 30 μg) was reverse transcribed using PrimeScript RT Master Mix (Takara). On-target or off-target sites of genomic DNA or cDNA were amplified with KOD Multi & Epi PCR kits (Toyobo) to generate sequencing libraries (see the Supplementary Table for the primer sequences). These libraries were sequenced using MiniSeq with a TruSeq HT Dual Index system (Illumina) as described before (41). Briefly, equal amounts of the PCR amplicons were subjected to paired-end sequencing using the Illumina MiniSeq platform. After MiniSeq, paired-end reads were analyzed by comparing wild type and mutant sequences using BE-analyzer (42).

### Quantitative real-time polymerase chain reaction (qRT-PCR)

The AAV copies in the RPE tissues were quantitatively estimated using the AAVpro Titration Kit (Takara) as previously described (7). The extracted AAVs from the RPE tissues underwent qRT-PCR according to the manufacturer’s instructions. The number of AAV genome copies was calculated using the standard curve with the Positive Control in the kit. The number of diploid mouse cells was estimated using a conversion factor of 1.6 × 10^4^ cells per 100 μg of genomic DNA.

### Immunofluorescence

At 6 weeks after subretinal injection, RPE-choroid-scleral complexes were prepared from enucleated eyes. The whole mount tissues were immunostained with antiHA antibody (1:100; cat. no. 26183-D680, Thermo) and anti-RPE65 antibody (1:100; cat. no. NB100-355AF488, Novus). Nuclear staining was performed using 4’,6-diamidino-2-phenylindole dihydrochloride (Sigma). Then, the slides were observed under a confocal microscope (Leica).

### ERG

Mice were dark-adapted overnight. After deep anesthesia, pupils were dilated by topical administration of phenylephrine hydrochloride (5 mg/ml) and tropicamide (5 mg/ml). Fullfield electroretinography was performed using the universal testing and electrophysiologic system 2000 (UTAS E-2000, LKC). The responses were recorded at a gain of 2 k utilizing a notch filter at 60 Hz and were bandpass filtered between 0.1 and 1500 Hz. The Prism 9 (GraphPad) was used to visualize the graphs and estimate the amplitudes. The amplitudes of the a-wave were measured from the baseline to the lowest negative-going voltage, whereas peak b-wave amplitudes were calculated from the trough of the a-wave to the highest peak of the positive b-wave.

### Optokinetic response

The virtual optokinetic system (OptoMotry HD, CerebralMechanics) was used to measure the grating acuity as visual thresholds to drive head tracking of mice to a rotating virtual cylinder, according to the manufacturer’s instructions and original publications on the system (43, 44). Briefly, mice were placed on a platform where they were exposed to view the rotating cylinder on the monitors. Visual thresholds were determined with a staircase procedure to produce the maximum spatial frequency (cycles/degrees) above which the mice did not respond to the rotating stimuli.

### Histologic evaluation

At 6 weeks after subretinal injection, paraffin blocks were prepared from enucleated eyes. After H&E staining, thin sections were evaluated for histologic toxicity. To investigate the histologic toxicity of adenine base editing, H&E-stained slides were evaluated to measure the ratio of the thickness of retinal layers (the thickness from the internal limiting membrane to the inner nuclear layer to that from the internal limiting membrane to the outer nuclear layer) (45).

### Statistics

All group results are expressed as mean ± SEM, if not stated otherwise. Comparisons between groups were made using one-way ANOVA and Tukey post-hoc multiple comparison tests. Statistical analyses were performed using Prism 9 (GraphPad).

### Study approval

All animal experiments were performed following the Association for Research in Vision and Ophthalmology statement for the use of animals in ophthalmic and vision research. The protocols were approved by the Institutional Animal Care and Use Committee of both Seoul National University.

## Author contributions

DHJ performed/analyzed animal experiments and wrote the manuscript. HJK, JHH, GR, and YJ performed/analyzed cell studies. CSC performed animal experiments. SB and JHK conceived/supervised this project and wrote the manuscript.

## Acknowledgments

This work was supported by the New Faculty Startup Fund from Seoul National University and by grants from the National Research Foundation of Korea (NRF) (no. 2017M3A9B4062654 and no. 2018M3D1A1058826 to J.H.K.; no. 2020M3A9I4036074 and no. 2017R1A6A3A04004741 to D.H.J.; and no. 2018M3A9H3022412 and no. 2020M3A9I4036072 to S.B.) and the Technology Innovation Program (no. 20000158 to S.B).

## References

1. Cideciyan AV. Leber congenital amaurosis due to RPE65 mutations and its treatment with gene therapy. Prog Retin Eye Res. 2010;29(5):398–427.

2. Sahel JA. Spotlight on childhood blindness. J Clin Invest. 2011;121(6):2145–9.

3. Cideciyan AV, Jacobson SG. Leber Congenital Amaurosis (LCA): Potential for Improvement of Vision. Invest Ophthalmol Vis Sci. 2019;60(5):1680–95.

4. Russell S, et al. Efficacy and safety of voretigene neparvovec (AAV2-hRPE65v2) in patients with RPE65-mediated inherited retinal dystrophy: a randomised, controlled, open-label, phase 3 trial. Lancet. 2017;390(10097):849–60.

5. DiCarlo JE, Mahajan VB, Tsang SH. Gene therapy and genome surgery in the retina. J Clin Invest. 2018;128(6):2177–88.

6. Maeder ML, et al. Development of a gene-editing approach to restore vision loss in Leber congenital amaurosis type 10. Nat Med. 2019;25(2):229–33.

7. Jo DH, et al. CRISPR-Cas9-mediated therapeutic editing of Rpe65 ameliorates the disease phenotypes in a mouse model of Leber congenital amaurosis. Sci Adv. 2019;5(10):eaax1210.

8. Jang HK, Song B, Hwang GH, Bae S. Current trends in gene recovery mediated by the CRISPR-Cas system. Exp Mol Med. 2020;52(7):1016–27.

9. Kosicki M, Tomberg K, Bradley A. Repair of double-strand breaks induced by CRISPR-Cas9 leads to large deletions and complex rearrangements. Nat Biotechnol. 2018;36(8):765–71.

10. Adikusuma F, et al. Large deletions induced by Cas9 cleavage. Nature. 2018;560(7717):E8–E9.

11. Rees HA, Liu DR. Base editing: precision chemistry on the genome and transcriptome of living cells. Nat Rev Genet. 2018;19(12):770–88.

12. Gaudelli NM, et al. Programmable base editing of A*T to G*C in genomic DNA without DNA cleavage. Nature. 2017;551(7681):464–71.

13. den Hollander AI, Roepman R, Koenekoop RK, Cremers FP. Leber congenital amaurosis: genes, proteins and disease mechanisms. Prog Retin Eye Res. 2008;27(4):391–419.

14. Astuti GD, et al. Comprehensive genotyping reveals RPE65 as the most frequently mutated gene in Leber congenital amaurosis in Denmark. Eur J Hum Genet. 2016;24(7):1071–9.

15. Pang JJ, et al. Retinal degeneration 12 (rd12): a new, spontaneously arising mouse model for human Leber congenital amaurosis (LCA). Mol Vis. 2005;11:152–62.

16. Jespersgaard C, et al. Molecular genetic analysis using targeted NGS analysis of 677 individuals with retinal dystrophy. Sci Rep. 2019;9(1):1219.

17. Zhong Z, et al. Seven novel variants expand the spectrum of RPE65-related Leber congenital amaurosis in the Chinese population. Mol Vis. 2019;25:204–14.

18. Jeong YK, Song B, Bae S. Current Status and Challenges of DNA Base Editing Tools. Mol Ther. 2020;28(9):1938–52.

19. Koblan LW, et al. Improving cytidine and adenine base editors by expression optimization and ancestral reconstruction. Nat Biotechnol. 2018;36(9):843–6.

20. Nishimasu H, et al. Engineered CRISPR-Cas9 nuclease with expanded targeting space. Science. 2018;361(6408):1259–62.

21. Jeong YK, Yu J, Bae S. Construction of non-canonical PAM-targeting adenosine base editors by restriction enzyme-free DNA cloning using CRISPR-Cas9. Sci Rep. 2019;9(1):4939.

22. Levy JM, et al. Cytosine and adenine base editing of the brain, liver, retina, heart and skeletal muscle of mice via adeno-associated viruses. Nat Biomed Eng. 2020;4(1):97–110.

23. Stevens AJ, Sekar G, Shah NH, Mostafavi AZ, Cowburn D, Muir TW. A promiscuous split intein with expanded protein engineering applications. Proc Natl Acad Sci U S A. 2017;114(32):8538–43.

24. Truong DJ, et al. Development of an intein-mediated split-Cas9 system for gene therapy. Nucleic Acids Res. 2015;43(13):6450–8.

25. Yu W, Wu Z. Ocular delivery of CRISPR/Cas genome editing components for treatment of eye diseases. Adv Drug Deliv Rev. 2020.

26. Suh S, et al. Restoration of visual function in adult mice with an inherited retinal disease via adenine base editing. Nat Biomed Eng. 2020. doi:10.1016/j.addr.2020.06.011

27. Redmond TM, et al. Rpe65 is necessary for production of 11-cis-vitamin A in the retinal visual cycle. Nat Genet. 1998;20(4):344–51.

28. Liang P, et al. Genome-wide profiling of adenine base editor specificity by EndoV-seq. Nat Commun. 2019;10(1):67.

29. Rees HA, Wilson C, Doman JL, Liu DR. Analysis and minimization of cellular RNA editing by DNA adenine base editors. Sci Adv. 2019;5(5):eaax5717.

30. Grünewald J, et al. Transcriptome-wide off-target RNA editing induced by CRISPR-guided DNA base editors. Nature. 2019;569(7756):433–7.

31. Zhou C, et al. Off-target RNA mutation induced by DNA base editing and its elimination by mutagenesis. Nature. 2019;571(7764):275–8.

32. Bae S, Park J, Kim JS. Cas-OFFinder: a fast and versatile algorithm that searches for potential off-target sites of Cas9 RNA-guided endonucleases. Bioinformatics. 2014;30(10):1473–5.

33. Tsai SQ, Nguyen NT, Malagon-Lopez J, Topkar VV, Aryee MJ, Joung JK. CIRCLE-seq: a highly sensitive in vitro screen for genome-wide CRISPR–Cas9 nuclease off-targets. Nat Methods. 2017;14(6):607–14.

34. Kim J, Malashkevich V, Roday S, Lisbin M, Schramm VL, Almo SC. Structural and kinetic characterization of Escherichia coli TadA, the wobble-specific tRNA deaminase. Biochemistry. 2006;45(20):6407–16.

35. Richter MF, et al. Phage-assisted evolution of an adenine base editor with improved Cas domain compatibility and activity. Nat Biotechnol. 2020;38(7):883–91.

36. Y. K. Jeong, et al. Nature Research. 2020. doi:10.21203/rs.3.rs-84912/v1

37. Gaudelli NM, et al. Directed evolution of adenine base editors with increased activity and therapeutic application. Nat Biotechnol. 2020;38(7):892–900.

38. Lee C, et al. CRISPR-Pass: Gene Rescue of Nonsense Mutations Using Adenine Base Editors. Mol Ther. 2019;27(8):1364–71.

39. Hug N, Longman D, Caceres JF. Mechanism and regulation of the nonsense-mediated decay pathway. Nucleic Acids Res. 2016;44(4):1483–95.

40. Park SW, Kim JH, Park WJ, Kim JH. Limbal Approach-Subretinal Injection of Viral Vectors for Gene Therapy in Mice Retinal Pigment Epithelium. J Vis Exp. 2015(102):e53030.

41. Bae S, Kweon J, Kim HS, Kim JS. Microhomology-based choice of Cas9 nuclease target sites. Nat Methods. 2014;11(7):705–6.

42. Hwang GH, et al. Web-based design and analysis tools for CRISPR base editing. BMC Bioinformatics. 2018;19(1):542.

43. Douglas RM, Alam NM, Silver BD, McGill TJ, Tschetter WW, Prusky GT. Independent visual threshold measurements in the two eyes of freely moving rats and mice using a virtual-reality optokinetic system. Vis Neurosci. 2005;22(5):677–84.

44. Prusky GT, Alam NM, Beekman S, Douglas RM. Rapid quantification of adult and developing mouse spatial vision using a virtual optomotor system. Invest Ophthalmol Vis Sci. 2004;45(12):4611–6.

45. Kim JH, et al. Absence of intravitreal bevacizumab-induced neuronal toxicity in the retina. Neurotoxicology. 2008;29(6): 1131–5.

