## Supplemental figures and tables for "Therapeutic adenine base editing corrects nonsense mutation and improves visual function in a mouse model of Leber congenital amaurosis"

**Table of Contents**

**Supplemental Figure 1.** The activity of dual ABE vectors.

**Supplemental Figure 2.** The five most common sequences in mRNA sequencing of the RPE cells from *rd12* mice treated with AAV-ABE.

**Supplemental Figure 3.** Histological analysis of retinal tissues at 6 weeks after the subretinal injection of AAV-ABE on wild-type C57BL/6 mice.

**Supplemental Table 1:** DNA off-target sites analyzed in this study.

**Supplemental Table 2:** RNA off-target sites analyzed in this study.

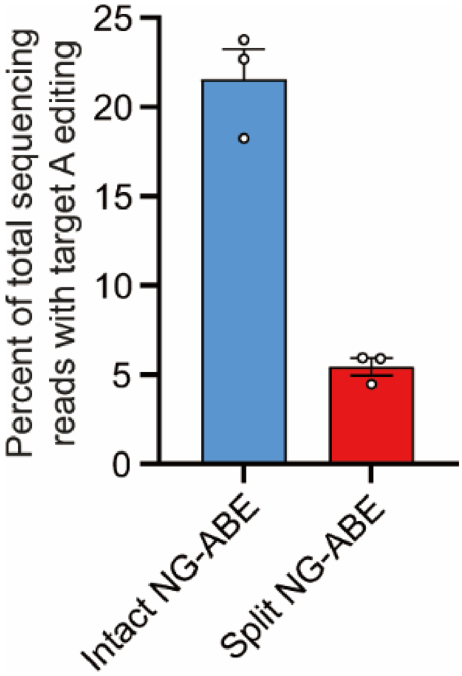


**Supplemental Figure 1. The activity of dual ABE vectors.** Quantitation of the intended A to G correction at position 6 (A6) in the mouse embryonic fibroblasts of *rd12* mice at 3 days after transfection of dual AAV encoded by plasmids.

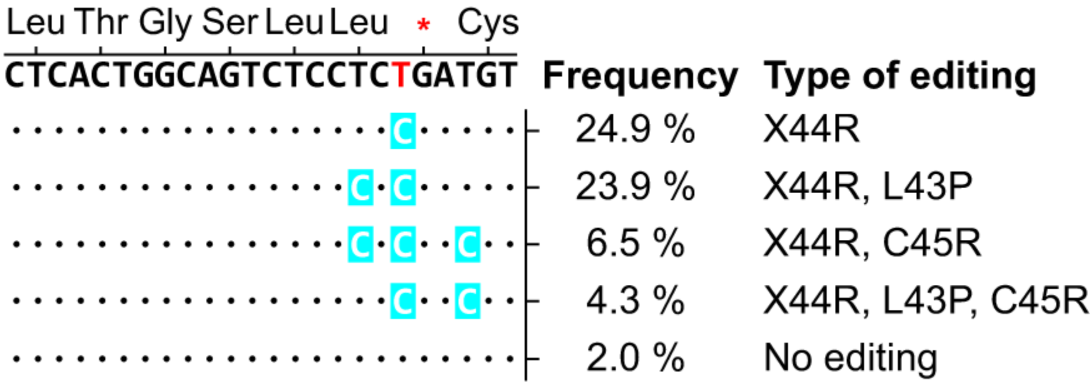


**Supplemental Figure 2. The five most common sequences in mRNA sequencing of the RPE cells from *rd12* mice treated with AAV-ABE.**

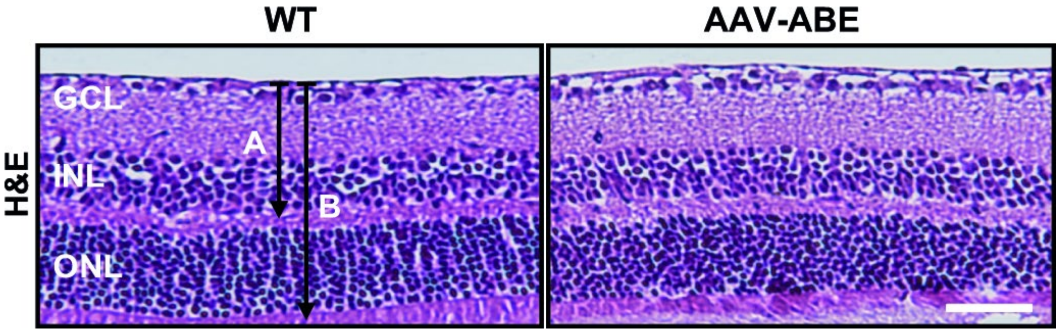


**Supplemental Figure 3. Histological analysis of retinal tissues at 6 weeks after the subretinal injection of AAV-ABE on wild-type C57BL/6 mice.** ‘A’ indicates the thickness from the internal limiting membrane to the inner nuclear layer and ‘B’ indicates that from the internal limiting membrane to the outer nuclear layer. GCL, ganglion cell layer; INL, inner nuclear layer; ONL, outer nuclear layer. Scale bar, 50 µm.

**Supplemental Table 1. DNA off-target sites analyzed in this study.** PAMs were distinguished from potential protospacer sequences by a tab. Mismatches to the on-target protospacer sequences were indicated with red upper cases. Bulges to the on-target protospacer sequences were indicated with red lower cases or hyphens. The position indicates potential cleavage sites (3 nt distal from PAM).

| Target | Sequence | Chromosome | Position |
|---|---|---|---|
| **rd12** | ACATCAGAGGAGACTGCCAG TGA | chr3 | 159601542 |
| **Offinder-1** | ACATCAGAGGAG-CTGACAG TGG | chrX | 117735664 |
| **Offinder-2** | ACAGCAGAGGAGACTGCtCAG TGG | chr3 | 22319361 |
| **Circle-1** | AAGTCAGAGGAGGCTGCCAG TGG | chr11 | 44913708 |
| **Circle-2** | TCATCAGAGGGAGATTGCCAG GGG | chr8 | 117198387 |
| **Circle-3** | A-ATCAGAGAACACTGCCAG AGG | chr2 | 67928117 |
| **Circle-4** | AGGTCAGAGGAGA-TGCCAG GGG | chr3 | 122497857 |
| **Circle-5** | GCATGAGAGGAGACTGACAG GTG | chr13 | 25042265 |
| **Circle-6** | TCAGCAGAGAAGACT-CCAG TGG | chr5 | 31469676 |
| **Circle-7** | GTATCAGAGAAGACTGCCAG AGG | chr4 | 63653115 |
| **Circle-8** | AAATGAGAGGAGACTCCAGG AGT | chr12 | 31448934 |
| **Circle-9** | TCATCAGAGAAG-CTGCCAG AGG | chr1 | 59436439 |
| **Circle-10** | AAGTCAGAGGAGGCTGCAGA GGG | chr3 | 148510669 |

**Supplemental Table 2. RNA off-target sites analyzed in this study.** Nucleotides that were predicted to be edited were shown in blue. The position indicates each A nucleotide.

| Gene | Sequence | Position |
|---|---|---|
| TadA Substrate | GCUCGGCUACGAACCGAG | tRNA from *E.coli* |
| Aars1 | GAUCCACUAUGA-CCGCA | chr20_34412828 |
| Perp | CGGCUCCUACGACGAUGG | chr10_18721122 |
| Flot1 | GAGGGACUAUGAGCUGAA | chr17_36136706 |
| MCM3AP | GGCAGCCUACGUAGCCGCA | chr10_76326644 |
| NCLN | CAUGGCGUACACAGCCGU | chr10_81323527 |
